# Sex-associated differences in cytomegalovirus prevention: Prophylactic strategy is associated with a strong kidney function impairment in female renal transplant patients

**DOI:** 10.1101/726968

**Authors:** Arturo Blazquez-Navarro, Chantip Dang-Heine, Chris Bauer, Nicole Wittenbrink, Kerstin Wolk, Robert Sabat, Oliver Witzke, Timm H. Westhoff, Birgit Sawitzki, Petra Reinke, Oliver Thomusch, Christian Hugo, Nina Babel, Michal Or-Guil

**Author notes:** Corresponding authors contact information Prof. Nina Babel: Berlin-Brandenburger Centrum für Regenerative Therapien, Charité-Universitätsmedizin Berlin, Augustenburger Platz 1, 13353 Berlin, Germany., Michal Or-Guil: Systems Immunology Lab, Department of Biology, Humboldt-Universität zu Berlin: Philippstr. 13, 10115 Berlin, Germany. These authors contributed equally (Nina Babel, Michal Or-Guil).

## Abstract

Post-transplantation cytomegalovirus (CMV) syndrome can be prevented using the antiviral drug (val)ganciclovir. (Val)ganciclovir is typically administered following a prophylactic or a pre-emptive strategy. The prophylactic strategy entails early universal administration, the pre-emptive strategy, early treatment in case of infection. However, it is not clear which strategy is superior with respect to transplantation outcome; sex-specific effects of these prevention strategies are not known. We have retrospectively analysed 540 patients from the multi-centre Harmony study along eight pre-defined visits: 308 were treated according to a prophylactic, 232 according to a pre-emptive strategy. As expected, we observed an association of prophylactic strategy with lower incidence of CMV syndrome, delayed onset and lower viral loads compared to the pre-emptive strategy. However, in female patients, the prophylactic strategy was associated with a strong impairment of glomerular filtration rate one year post-transplant (difference: -12.0±4.2 mL·min^-1^·1.73m^-2^, P=0.005). Additionally, we observed a tendency of higher incidence of acute rejection and severe BK virus reactivation in the prophylactic strategy group. While the prophylactic strategy was more effective for preventing CMV syndrome, our results suggest for the first time that the prophylactic strategy might lead to inferior transplantation outcomes in female patients, providing evidence for a strong association with sex.

## 1. Introduction

Cytomegalovirus (CMV) is a herpes virus often reported as the most important viral pathogen after kidney transplantation.^1–3^ It is a major cause of morbidity and mortality, being associated with retinitis, pneumonitis, colitis, encephalitis, allograft damage and allograft loss, among others.^1–4^ CMV syndrome or disease may occur as a consequence of reactivation of latent infections or through primary infection, acquired from the donor or from the environment.^2^ The major risk factor for CMV syndrome or disease is the pre-transplantation serostatus: CMV seronegative transplant recipients with a seropositive donor (D^+^R^-^) have the highest risk, while seropositive recipients (R^+^) have an intermediate risk and seronegative recipients with seronegative donors (D^-^R^-^) have the lowest risk.^2^ Moreover, the use of immunosuppressive drugs like rabbit antithymocyte globulin (ATG) can additionally increase the incidence of CMV (re)activations.^5^

The standards in prevention and treatment of CMV (re)activation are based on ganciclovir or its oral prodrug valganciclovir.^6,7^ In addition to antiviral therapy, CMV-specific T cell immunity has been shown to control CMV viral reactivations, determining the outcome of disease.^8–10^ Two prevention strategies are routinely employed in the clinic: prophylactic and pre-emptive.^2,6,7^ The prophylactic strategy is based on the universal administration of (val)ganciclovir in case of patients with a CMV risk constellation, usually during 3-6 months after transplantation.^6,7^ In the pre-emptive strategy, patients are regularly monitored for CMV through quantitative polymerase chain reaction or pp65 antigenemia test; (val)ganciclovir is only administered after a positive test, ideally before any symptoms of CMV syndrome or disease manifest.^6,7^ The pre-emptive strategy thus leads to a reduction of unnecessary treatments, which is advantageous with respect to the appearance of side effects and resistances against antiviral drugs.^6,7^

While the KDIGO guideline of 2009 preferred prophylaxis as the standard of prevention, the more recent reference CMV management guideline recommends both strategies for the prevention of CMV disease in patients with both high or intermediate CMV mismatch-based risk constellation.^6,7^ However, the differences in outcome with regard to other criteria, including renal function and other viral (re)activations is largely unclear. Interestingly, there is evidence of sex differences in both ganciclovir pharmacokinetics and the anti-CMV immune response.^11–18^ Thus, female patients have been shown to have a faster ganciclovir clearance, and distinct anti-CMV immunological profiles e.g. higher number of secreting anti-CMV T cells.^11–14,18^ In spite of this, there are to our knowledge still no studies on the influence of sex on the clinical outcomes of CMV prevention strategies. In this work, we provide evidence that prophylaxis might be associated with inferior transplantation outcomes in female patients.

## 2. Materials and Methods

### 2.1. Patient population

As part of the systems medicine project e:KID, we conducted a sub-study within the randomized, multi-centre, investigator-initiated Harmony trial (NCT 00724022)^19,20^ to determine the impact of CMV prevention strategy on transplant outcome. For this, CMV, Epstein-Barr virus (EBV) and BK virus (BKV) viral loads, white blood cell count and creatinine were measured at predetermined eight study visits.^20^ This viral monitoring was non-interventional and centrally performed and was independent from the internal, interventional viral monitoring (see section 2.3). The study was carried out in compliance with the Declaration of Helsinki and Good Clinical Practice.

### 2.2. Patient medication

According to study design, patients were treated with a quadruple (arm A) or triple (arms B and C) immunosuppressive therapy.^19^ Patients in arm A received an induction therapy with basiliximab and maintenance therapy consisting of tacrolimus (Advagraf®, Astellas), mycophenolate mofetil (MMF) and corticosteroids. Patients in arm B received the same treatment as in arm A, but corticosteroids were withdrawn at day 8. Patients in arm C received the same treatment as in arm B, except induction was achieved with ATG, instead of basiliximab.

### 2.3. Patient monitoring

Patients were monitored for transplantation outcomes during the first post-transplantation year. Graft function was monitored along eight visits, scheduled at day 0 (pre-transplantation), 2^nd^ week, 1^st^ month, 2^nd^ month, 3^rd^ month, 6^th^ month, 9^th^ month, and 12^th^ month. To assess graft function, glomerular filtration rate was calculated using the CKD-EPI formula, measured in mL·min^-1^·1.73 m^-2^.^21^ Serious adverse events were defined following the Good Clinical Practice guidelines. Suspected episodes of acute rejection had to be confirmed through biopsy; histologic characteristics were described according to the Banff criteria of 2005.^22^ Regarding the outcome assessment, acute rejection was analysed excluding borderline rejections. Routine surveillance biopsies were allowed but not mandatory.

### 2.4. Clinical monitoring and management of clinical complications

Viral (re)activations were monitored during the first post-transplantation year and managed at local centres as described previously.^20^ CMV in particular was monitored for all patients, independently of the prevention strategy. Monitoring was performed independently from the above described CMV viral load measurements and was based on three different methods: serum PCR viral load measurements; test for pp65 antigenemia and symptom monitoring according to the internal centre standards. Diagnosis of CMV syndrome was likewise based on the methods, where a qPCR over 1000 copies·mL^-1^ was defined as positive. Patients with CMV syndrome were treated based on internal centre standards. Suggested treatment was (val)ganciclovir treatment according to local standards with/or without reduction of tacrolimus and MMF dose. No data on the time point of CMV syndrome diagnostic were available for this study; no data on CMV disease were available.

### 2.5. Screening of CMV, EBV and BKV viraemia

In parallel to the clinical monitoring performed at each centre, peripheral blood samples from the eight visits were centrally monitored for CMV, EBV and BKV by TaqMan qPCR, as described previously.^20^ The centralized viral load assessment was non-interventional.

### 2.6. Definition of CMV prevention strategy groups and characterization of antiviral treatments

Patients were stratified into two prevention strategy groups based on the (val)ganciclovir treatments during the first 14 days. All patients that started a (val)ganciclovir treatment during the first 14 days were assigned into the prophylactic strategy group; the rest of the sub-cohort was classified in the pre-emptive strategy group. The 14 day threshold was chosen to allow comparability with our previous prospective study on the topic (VIPP), in which recruiting took place during the first two post-transplant weeks.^23,24^ CMV syndrome was treated equally for both strategy groups, as explained above. Antiviral treatments with no data on the end time point were disregarded for the calculation of the treatment duration but considered for the calculation of median dose. Accordingly, reported MMF dose and tacrolimus trough correspond to the 14 day threshold.

### 2.7. Viraemia-based patient classification

To assess the efficacy of prevention strategies regarding viral (re)activations, patients were classified based on their peak viral load values for CMV, EBV and BKV, as previously published.^20^ Briefly, the classifications are defined as follows: “detectable viral load” corresponds to patients with at least one viral load measurement over detection limit (250 copies·mL^-1^)^20^, “elevated viral load” to patients with at least one viral load measurement over 2000 copies·mL^-1^, “high viral load” to patients with at least one viral load measurement over 10000 copies·mL^-1^. These groups can overlap with each other.

### 2.8. Descriptive statistics and baseline analysis

Categorical variables are summarised here as numbers and frequencies; quantitative variables are reported as median and interquartile range (IQR). Differences between the groups were calculated using Pearson’s chi-square test with continuity correction (or two-tailed Fisher’s exact test, when stated); odds ratios (OR) and 95% confidence intervals (95% CI) are provided. In all cases, odds ratio over 1 denote a higher prevalence of the adverse event in the pre-emptive strategy group. Differences in quantitative variables between groups were analysed using the two-tailed Mann-Whitney test. Kaplan-Meier curves for time to occurrence of the first CMV (re)activation were calculated using the R survival package (version 2.43-3); strategy groups were compared using the log-rank test. Correlations are reported employing Spearman’s rho and P value). Box plots depict the median, first and third quartile of a variable; the maximum length of the whiskers corresponds to 1.5 times the IQR.

In baseline analysis, a P value below 0.050 was considered significant. For descriptive statistics, P values are reported purely for illustrative reasons – no definition of statistical significance is employed.

### 2.9. Multi-parameter regression modelling

To determine the influence of prevention strategies, sex and their interaction on transplantation outcomes, we performed multi-parameter regression controlling for confounders: The regression models used for the analysis incorporate as independent variables the prevention strategies, sex and their interaction term, as well as all selected confounding factors, and as dependent variable the outcome of interest. For categorical binary outcomes, logistic regression was employed, for continuous outcomes linear regression was used. The choice of confounders was performed through backward elimination by Akaike’s information criterion starting from a full model, as the criteria for the allocation to a prevention strategy are unknown so that a selection based on medical criteria is not possible. The full model incorporated – apart from prevention strategies, sex and their interaction – all measured demographic factors (see Table 1 and Table S1) and the transplantation centre. For the analysis of the eGFR one year after transplantation (eGFR-1y), CMV, BKV and EBV peak viral loads and acute rejection were additionally included as potential confounders, as these events preceded or were simultaneous to eGFR-1y and might hence have an influence on it. Peak viral loads were included in the models (both as independent or dependent variables) as log-transformed (base 10); viral loads below detection level were set to zero. After backward elimination, the resulting model for each outcome was tested for multi-collinearity and (in the case of linear regression) for homoscedasticity: Multi-collinearity was assessed calculating the generalized variance-inflation factor, with a threshold of 5 to exclude a factor. Homoscedasticity was evaluated with the studentized Breusch-Pagan test; if it cannot be assumed (P<0.050), robust standard errors are reported. The resulting model for each outcome is provided in the Supplementary Table S2.

**Table 1.** Differences in patient baseline characteristics between strategy groups. Data are given as number (percentage) or median [interquartile range]. Expanded criteria donors are defined as follows: age over 60 years or age over 50 years and at least two of the following factors: cerebrovascular accident as the cause of death, hypertension or a serum creatinine level over 1.5 mg·dL^-1^. P value is calculated based on Pearson’s chi-square test or Fisher’s exact test for binary variables (marked with ^a^) and based on Mann-Whitney test for continuous variables. Data on the cause of end-stage kidney disease are summarized in Table S1.

| Variable |  | Prophylactic strategy group (N=308) | Pre-emptive strategy group (N=232) | P value |
| --- | --- | --- | --- | --- |
| Female sex |  | 104 (33.8%) | 90 (38.8%) | 0.265 |
| Caucasian race |  | 304 (98.7%) | 231 (99.6%) | 0.397<sup>a</sup> |
| Recipient age (years) |  | 55 [46-64] | 57 [44-64] | 0.988 |
| Body mass index ($\text{kg}\cdot\text{m}^{-2}$) | | 26.3 [23.5-29.7] | 25.4 [22.8-28.4] | 0.059 |
| CMV mismatch-based risk | High (D<sup>+</sup>R<sup>-</sup>) | 119 (39.1%) | 27 (12.1%) | <0.001 |
|  | Medium (R<sup>+</sup>) | 137 (45.1%) | 129 (57.8%) |  |
|  | Low (D<sup>-</sup>R<sup>-</sup>) | 48 (15.8%) | 67 (30.0%) |  |
| EBV mismatch-based risk | High (D<sup>+</sup>R<sup>-</sup>) | 13 (5.1%) | 11 (6.2%) | 0.583<sup>a</sup> |
|  | Medium (R<sup>+</sup>) | 239 (93.4%) | 161 (91.0%) |  |
|  | Low (D<sup>-</sup>R<sup>-</sup>) | 4 (1.6%) | 5 (2.8%) |  |
| Donor age (years) |  | 55 [48-65] | 55 [46-65] | 0.931 |
| No previous transplantations |  | 298 (96.8%) | 216 (94.7%) | 0.346 |
| Living donor |  | 31 (10.1%) | 35 (15.4%) | 0.088 |
| Expanded criteria donor |  | 136 (44.2%) | 99 (42.7%) | 0.798 |
| Cold ischaemia time (min) |  | 626 [427-844] | 600 [414-840] | 0.505 |
| Number of HLA A, B and DR mismatches |  | 3 [2-4] | 3 [1-4] | 0.457 |
| No panel-reactive antibodies before transplantation |  | 23 (7.6%) | 17 (7.7%) | 1.000 |
| White blood cell count (cells·L<sup>-1</sup>) |  | 7.2 [5.7-8.9] | 7.1 [6.0-8.5] | 0.676 |
| Therapy arm | A (basiliximab+steroids) | 93 (30.2%) | 96 (41.4%) | <0.001 |
|  | B (basiliximab) | 92 (29.9%) | 83 (35.8%) |  |
|  | C (ATG) | 123 (39.9%) | 53 (22.8%) |  |
| Low MMF daily dose (< 2000 mg·day<sup>-1</sup>) |  | 37 (12%) | 45 (19.4%) | 0.025 |
| Tacrolimus trough level (ng·mL<sup>-1</sup>) |  | 9.9 [7.4-12.5] | 9.2 [7-12] | 0.145 |

In the multi-parameter analysis, a P value below 0.050 was considered significant. P values were not corrected for multiple testing, as this study was of exploratory nature.^25–27^

## 3. Results

### 3.1. Definition of study sub-cohorts

To assess the effects of CMV prevention strategy on transplantation outcome, we retrospectively analysed the cohort of an existent study (N=540 patients from 18 centres) with a female ratio of 35.9% (N=194).^19,20^ Patients were grouped into two sub-cohorts, based on whether they started an antiviral therapy during the first two post-transplant weeks (prophylactic strategy group, N=308) or not (pre-emptive strategy group, N=232) (see 2.6). As described previously, viral load (CMV, EBV and BKV), graft function and other clinical markers were collected along eight visits during the first post-transplant year; a total of 3715 blood samples were analysed.^20^

In this work, we have evaluated the effects of prevention strategy and sex on the main outcome eGFR-1y and the secondary outcomes, incidence of acute rejection, CMV complications, and BKV and EBV (re)activations. The analyses were performed based on the following approach: For descriptive purposes, single-parameter differences between sub-cohorts were assessed; multi-parameter regression analysis – controlling for all potential confounders – was employed to determine any effects of prevention strategy, sex and their interaction on transplantation outcomes. After an assessment of the baseline characteristics of the sub-cohorts, we describe in detail the most important findings in the next sections.

### 3.2. Study sub-cohorts characteristics

To identify differences at baseline between the two prevention strategy sub-cohorts regarding demographics or treatment procedures, we performed comparative statistics (see Table 1, for cause of end-stage kidney disease see Table S1). As shown in Table 1, significant (P<0.050) differences were found for MMF daily dose, CMV mismatch-based risk and therapy arm; the difference was highly significant (P<0.001) for the latter two factors; specifically among female patients a significant difference in body mass index was found, but not in therapy arm (see Table S3).

59 patients (25.4%) of the pre-emptive strategy group were treated with (val)ganciclovir after the second post-transplantation week. In total, 367 patients (68.0%) received (val)ganciclovir during the first post-transplant year, independently of their prevention strategy group; use of antivirals in both groups is shown in Table 2.

**Table 2.** Antiviral treatment details for the two strategy groups. Data are given as number (percentage) or median [interquartile range].

| **Variable** | **Prophylactic**<br>**strategy group**<br>**(N=308)** | **Pre-emptive strategy**<br>**group treated with**<br>**(val)ganciclovir**<br>**(N=59)** |
| --- | --- | --- |
| Median time under<br>(val)ganciclovir (days) | 118 [87-182] | 92 [46-155] |
| Valganciclovir<br>average daily dose<br>(mg·day<sup>-1</sup>) | 277 [165-450] | 450 [205-454] |
| Ganciclovir usage | 34 (11.0%) | 8 (13.6%) |
| Intravenous<br>ganciclovir usage | 31 (10.1%) | 7 (11.9%) |

Regarding differences in outcomes between female and male patients, we observed no differences between sexes with respect (Table 3).

**Table 3.**
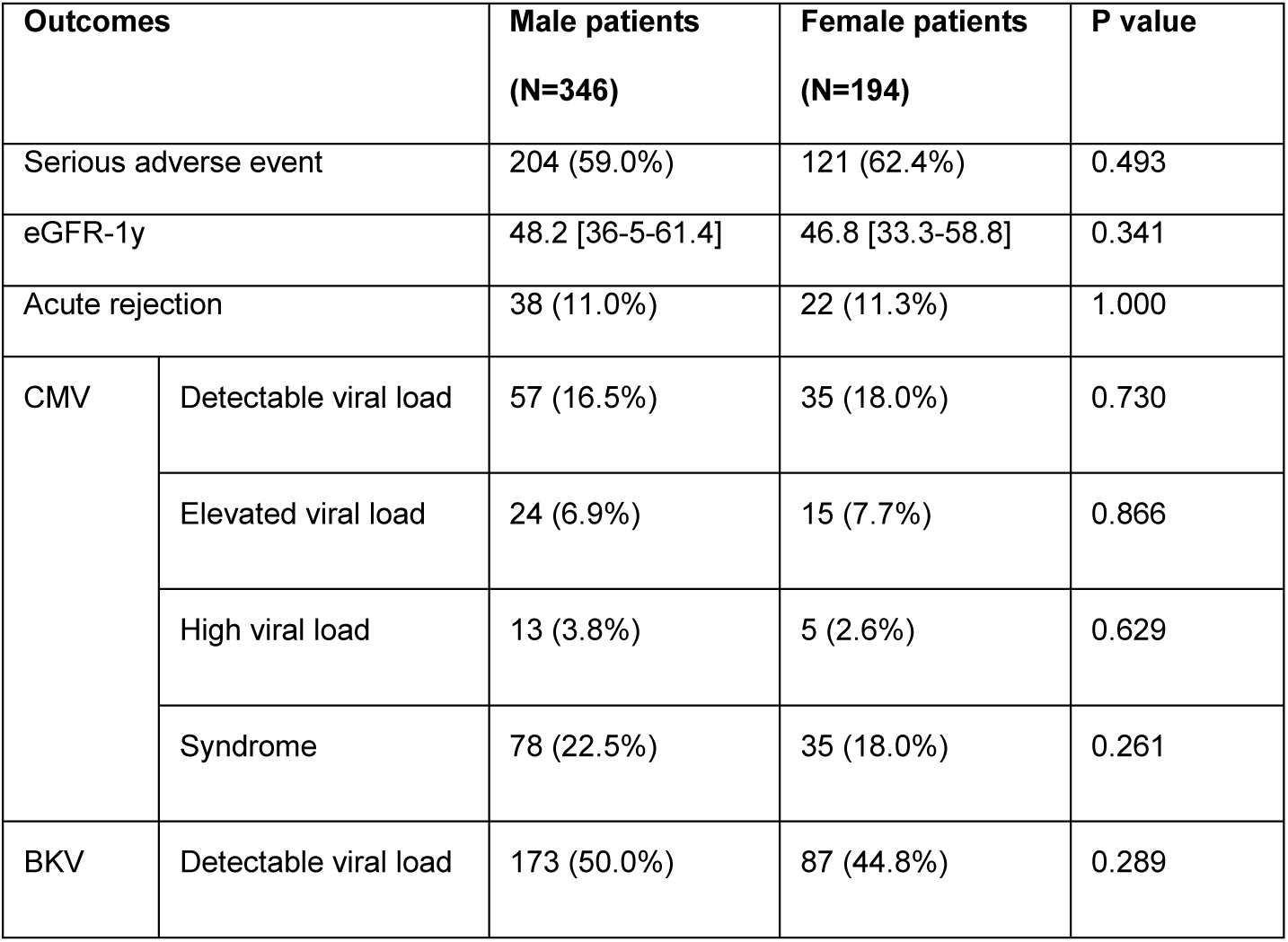
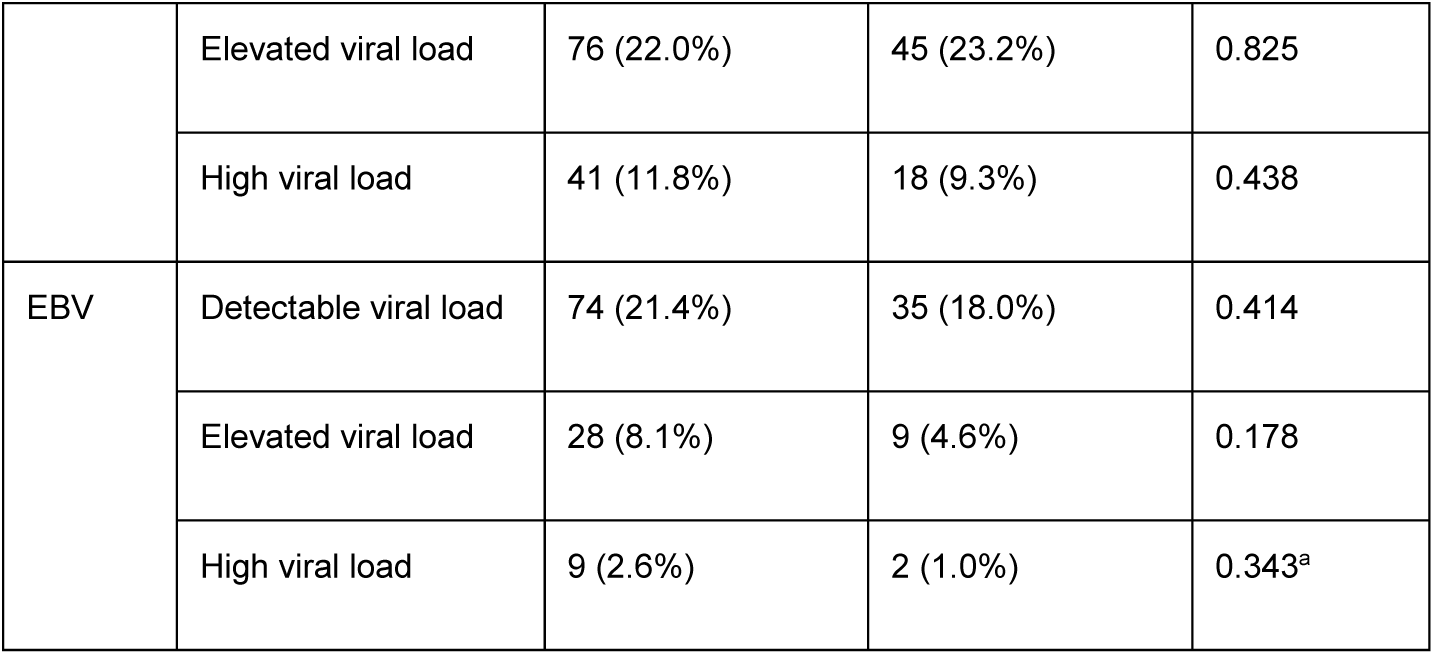
Differences between sexes in outcomes of the first post-transplantation year. Data are given as number (percentage) or median [interquartile range]. P value is calculated based on Pearson’s chi-square test or Fisher’s exact test for binary variables (marked with ^a^) and on Mann-Whitney test for continuous variables. For the definition of (re)activation severity degrees see Methods (2.7). As it can be observed, there were no differences between sexes with respect to the measured outcomes.

| **Outcomes** |  | **Male patients**<br>**(N=346)** | **Female patients**<br>**(N=194)** | **P value** |
| --- | --- | --- | --- | --- |
| Serious adverse event |  | 204 (59.0%) | 121 (62.4%) | 0.493 |
| eGFR-1y |  | 48.2 [36-5-61.4] | 46.8 [33.3-58.8] | 0.341 |
| Acute rejection |  | 38 (11.0%) | 22 (11.3%) | 1.000 |
| CMV | Detectable viral load | 57 (16.5%) | 35 (18.0%) | 0.730 |
|  | Elevated viral load | 24 (6.9%) | 15 (7.7%) | 0.866 |
|  | High viral load | 13 (3.8%) | 5 (2.6%) | 0.629 |
|  | Syndrome | 78 (22.5%) | 35 (18.0%) | 0.261 |
| BKV | Detectable viral load | 173 (50.0%) | 87 (44.8%) | 0.289 |
|  | Elevated viral load | 76 (22.0%) | 45 (23.2%) | 0.825 |
|  | High viral load | 41 (11.8%) | 18 (9.3%) | 0.438 |
| EBV | Detectable viral load | 74 (21.4%) | 35 (18.0%) | 0.414 |
|  | Elevated viral load | 28 (8.1%) | 9 (4.6%) | 0.178 |
|  | High viral load | 9 (2.6%) | 2 (1.0%) | 0.343<sup>a</sup> |

### 3.3. Prophylactic strategy group was associated with a serious impairment of graft function in female patients

A descriptive analysis showed that patients in the prophylactic strategy group had, in general, a poorer transplantation course than those in the pre-emptive strategy group, with a higher incidence of total serious adverse events (64.6% vs. 54.3%, P=0.020, OR: 0.65 [0.45-0.94]). For the main outcome, renal function, single-parameter analysis likewise revealed a difference between the prevention groups. Thus, eGFR-1y was lower in the prophylactic strategy group compared to the pre-emptive group (45.6 [33.5-58.3] vs. 50.3 [38.1-64.5] mL·min^-1^·1.73m^-2^, P=0.011). Of note, the difference in eGFR was noticeable for all visits from the third post-transplant month on (Figure S4).

Importantly, the impairment of eGFR-1y in the prophylactic group was only observed for female patients, with a difference of 18.5 mL·min^-1^·1.73m^-2^ (38.4 [28.8-53.6] vs. 56.8 [41.3-67.9] mL·min^-1^·1.73m^-2^, P<0.001). Among male patients, the prophylactic strategy group had a slightly higher median eGFR-1y (48.5 [36.3-61.5] vs. 47.2 [37.2-59.6] mL·min^-1^·1.73m^-2^, P=1.000). A difference in eGFR for females could be observed already one month after transplantation (Figure 1).

**Figure 1.**
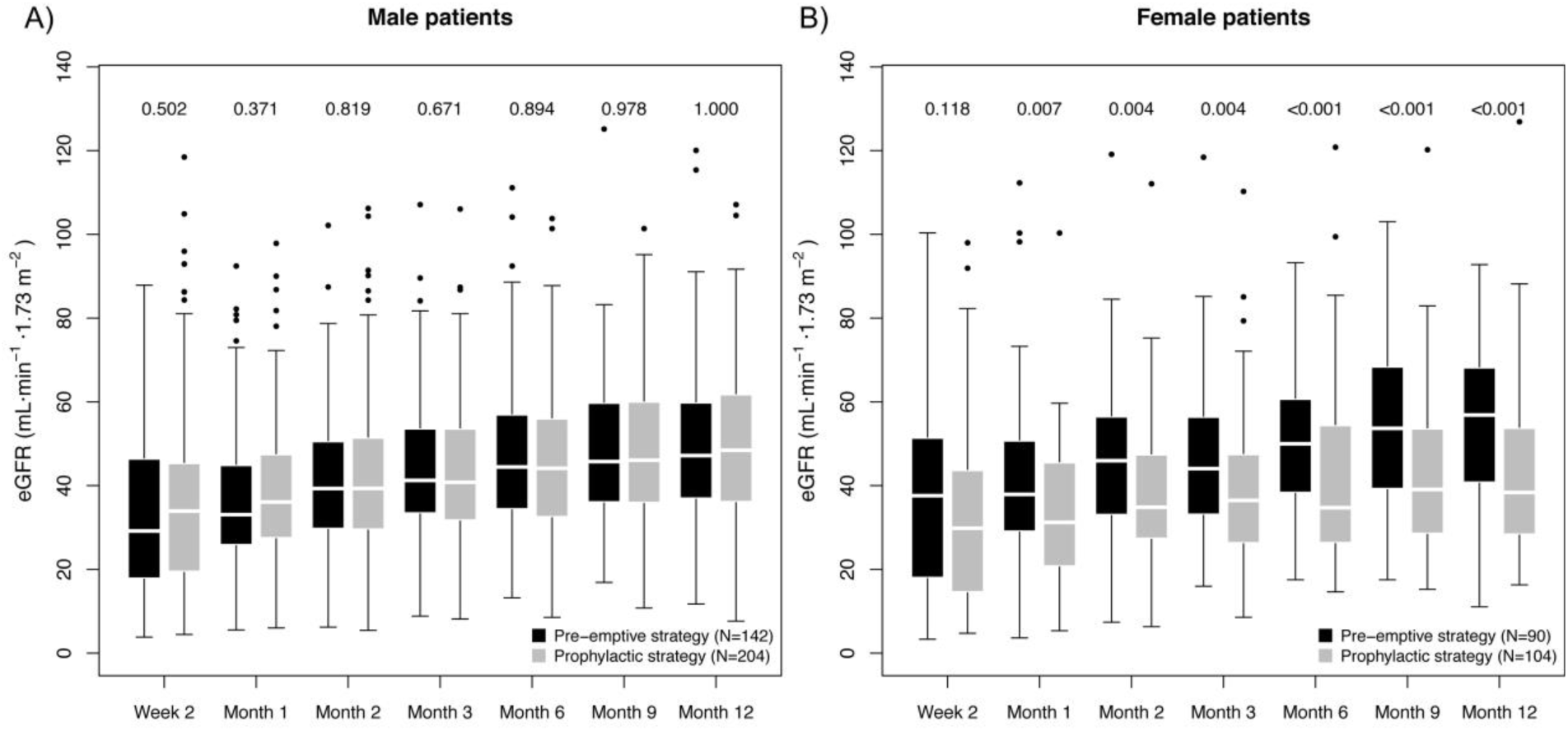
Box plot of the graft function dynamics of the prevention strategy groups stratified for sex. The numbers indicate the p value of the difference in eGFR between the prevention strategy groups, as calculated using the Mann-Whitney test. The P values for the first six measurements are included only to facilitate understanding on the eGFR dynamics, and are therefore not adjusted for multiple testing.

Multi-parameter regression incorporating all potential confounders confirmed a significant, strong association of the interaction term prophylactic strategy:female sex with decreased eGFR-1y (estimate: -12.0±4.2 mL·min^-1^·1.73m^-2^, P=0.005); while no significant association was found for prevention strategy (P=0.658) or sex alone (P=0.145). For more details on the multi-parameter regression model, see Table S2A. As a tendency towards lower eGFR in females under the prophylactic strategy (see Figure 1B) was already observed two weeks after transplantation (eGFR-2w), we tested additionally the possibility that a difference in baseline conditions as the cause for the observed association. Therefore, we repeated the analysis incorporating eGFR-2w as a confounder (Table S2B). Remarkably, in spite of the highly significant correlation of eGFR-2w with eGFR-1y (0.46±0.04 mL·min^-1^·1.73m^-2^, P<0.001), the negative effect of prophylactic strategy:female sex was consistently strong (−10.2±3.6 mL·min^-1^·1.73m^-2^, P=0.005). In conclusion, we did not find any evidence of spuriousness of the observed effect of prophylactic strategy:female sex on the eGFR-1y.

We further investigated the nature of the difference in eGFR between prevention strategies in female patients, examining the associations of dose and beginning of therapy with eGFR-1y. We did not observe any negative effect of high doses: We compared the female patients in the pre-emptive strategy group that received a valganciclovir treatment, with those in the prophylactic strategy group, as the first group had a higher daily dose than the second (P=0.041). Thus, we observed that these patients had a higher eGFR-1y than those in the prophylactic group (38.4 [28.8-53.6] vs. 57.7 [40.1-66.6] mL·min^-1^·1.73m^-2^, P=0.005), in spite of the higher valganciclovir dose. On the other hand, we observed an effect of therapy timing in eGFR-1y, with a positive correlation between day of treatment beginning and eGFR-1y (ρ=0.27, P=0.015) among female patients who received (val)ganciclovir.

### 3.4. The prophylactic strategy was associated with significantly lower CMV viral loads and incidence of syndrome than the pre-emptive strategy

We further evaluated the effectivity of the strategies in the prevention of CMV complications. The single-parameter, descriptive analysis showed a higher incidence of CMV viral load in the pre-emptive strategy group (19.8% vs. 14.9%, P=0.167); for CMV syndrome a higher incidence was found in the prophylactic strategy group (see Table S5 A). The latter was not unexpected, as most patients with high CMV risk were in the prophylactic strategy group (see Table 2). Stratifying for CMV risk, a clear trend for lower incidence of CMV (re)activation was observed in the prophylactic strategy group; but not for CMV syndrome (Table S5 B). However, the results of the multi-parameter regression (Tables S2 C-D) show that prophylactic strategy had a significant association with both lower peak CMV viral load (−0.73±0.23 log_10_(copies·mL^-1^), P=0.002) and CMV syndrome incidence (−1.17±0.57, P=0.042). No significant sex effects, nor interactions between prevention strategy and sex were observed for these outcomes.

Interestingly, CMV incidence showed different temporal patterns in the two strategy groups (Figure 2): While in the pre-emptive strategy group 86.7% of all CMV load events occurred in the first 100 days post-transplant, in the prophylactic strategy group it was only 56.1% (Figure 2A). Moreover, a higher prevalence of detectable CMV viral load was observed in the prophylactic strategy group for all study visits after the third month (Figure 2B).

**Figure 2.**
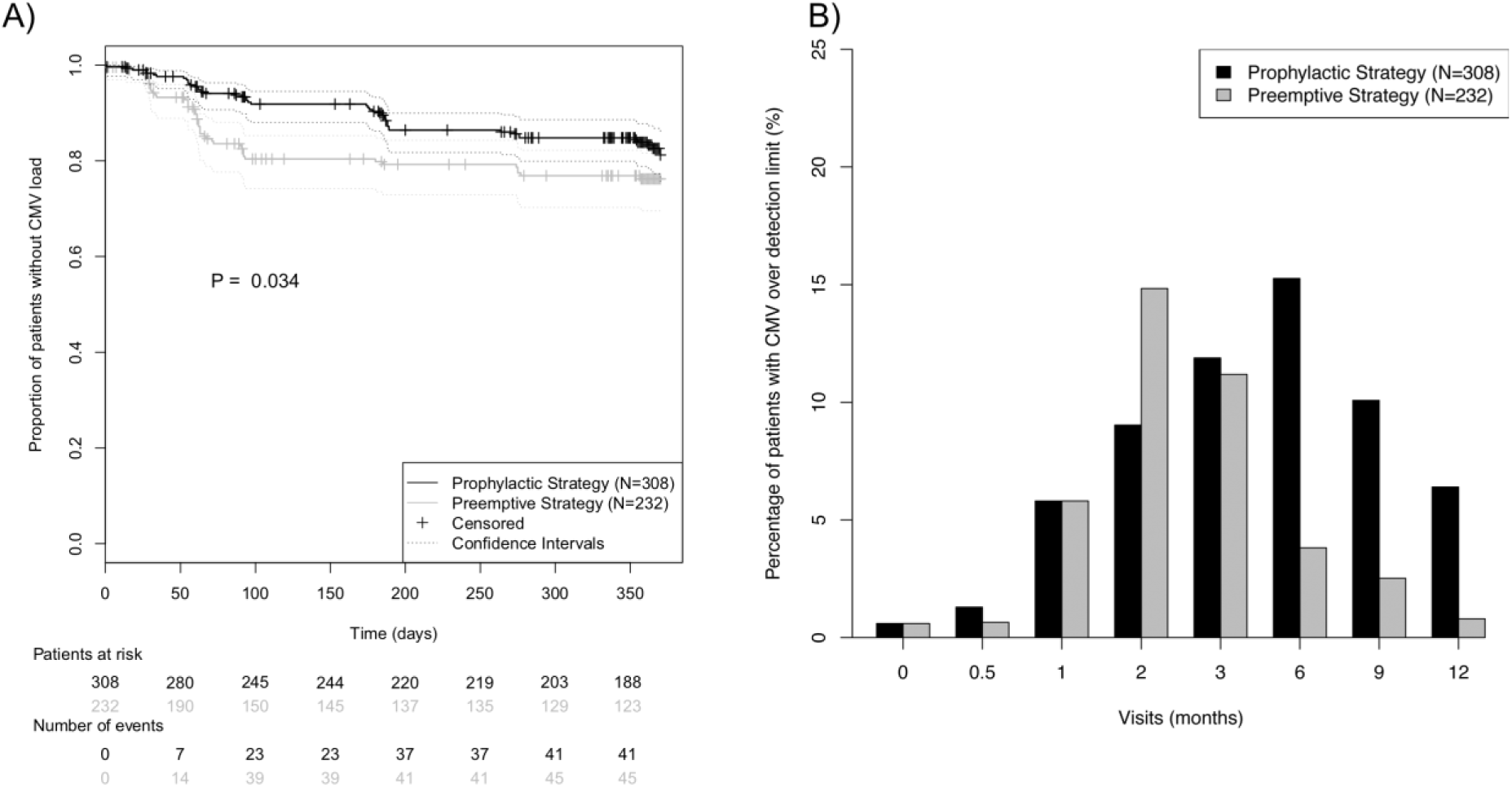
Incidence of CMV (re)activation in the prevention strategy groups during the first post-transplant year. (A) Kaplan-Meier curves for absence of CMV (re)activation during the first post-transplant year. CMV (re)activation was defined as viral load over detection limit. Prevention strategy groups were compared using the log-rank test. (B) Prevalence of CMV viral load over detection limit for each of the eight protocol visits.

### 3.5. There was a tendency towards higher incidence of acute rejection and BKV (re)activation among patients under prophylactic strategy

Regarding the important complication acute rejection, a tendency towards higher incidence was found in the prophylactic strategy group (14.9% vs. 6.0%, P=0.002, OR: 0.37 [0.18-0.70]). However, the multi-parameter analysis (Table S2E) could not confirm this tendency, as only a borderline significant association of prophylactic strategy group with rejection (−0.92±0.50, P=0.068) was observed; there was no evidence of an effect of sex, nor of the interaction with prevention strategy.

Regarding the effects of prevention strategy for other viruses, no evidence of an effect of prevention strategy was found for EBV, neither through (stratified) single-parameter analysis (Table S5 A and S5 C), nor through multi-parameter analysis (Table S2F). Furthermore, there was no evidence of sex-specific effects. On the other hand, we found a higher incidence of severe BKV (re)activation in patients of the prophylactic strategy group (P=0.056, OR: 0.55 [0.29-1.01]), see Table S5 A. The multi-parameter analysis (Table S2 G) showed likewise a borderline significant association of prophylactic strategy with higher BKV viral loads (0.55±0.29 log_10_(copies·mL^-1^), P=0.055), while no effect of sex could be observed.

## 4. Discussion

The goal of our study was to evaluate the clinical efficacy and sex-associated differences of two common CMV prevention strategies in a large cohort of kidney transplant patients from the multi-centre Harmony study. The main finding of the study is the first evidence of superiority of the pre-emptive strategy in female patients with respect to graft function.^23,24,28–34^ The observed effect was very large, corresponding to a increase of 12.0±4.2 mL·min ^-1^·1.73m^-2^ in eGFR-1y, which is especially relevant, as eGFR-1y is an accepted marker for long term transplantation outcomes.^35^ Even after controlling for differences in eGFR observed two weeks after transplantation (at a time point in which effects of prevention strategy are already thinkable), the effect of prevention strategy among female patients remained consistently strong.

Our results highlight the importance of sex-associated effects in transplantation. In recent years, sex differences have emerged as an essential factor in clinical studies.^36^ In transplantation, several complications are associated with sex, including acute rejection, graft loss and viral (re)activations.^11,13,37,38^ However, the underlying reasons for these sex differences are not well understood; possible causes include the hormonal regulation of the immune system, the effects of pregnancy, and differences in the metabolism of drugs routinely employed in transplantation.^11^ For example, there is tentative evidence of sex-related differences in the pharmacokinetics of (val)ganciclovir.^11,12^ Thus, ganciclovir clearance has been observed to be 24% faster in female transplantation patients, suggesting higher activity of the organic anion transporter 1.^11,12^ Furthermore, it has been shown repeatedly that women and men have different anti-CMV immunological profiles, with an impact in the graft function and even the phenotype of the immune system as a whole.^13,14,18^ Notably, in a recent publication, Lindemann *et al.* have observed an association of high numbers of IL-21-secreting anti-CMV T cells with female sex and lower eGFR in a clinical transplantation context.^13^

Our analyses may provide some evidence on the nature of the observed association of eGFR-1y with prevention strategies in female patients. Although the impaired graft function in the female prophylactic strategy group can be partly explained through the higher incidence of BKV severe (re)activation and rejection, the results of the multi-parameter analysis showed an independent association of prevention strategy with graft function, regardless of these adverse events.^20,39,40^ Therefore, our results do not support the hypothesis that these adverse events are the main cause for the difference in eGFR-1y between sub-cohorts. Regarding possible nephrotoxic effects of the antiviral drug, we did not find any association of higher valganciclovir doses with lower eGFR – rather, the opposite association was observed – in contrast to Heldenbrand *et al*.^41^ The absence of a negative dose-dependent effect suggests that the observed difference was not a consequence of nephrotoxicity of valganciclovir. On the other hand, the time of beginning of the (val)ganciclovir therapy was determinant for the eGFR-1y: the later patients began the therapy, the higher the renal function.

Albeit being highly speculative, we hypothesize that the observed results may be (at least in part) caused by an immunological mechanism. As we previously demonstrated, an increased number of CMV-specific T-cells upon CMV (re)activation is associated with reduced alloreactivity and improved graft function in renal transplantation patients.^42^ Similarly, in liver transplantation, primary CMV infection has been found to be associated with donor-specific CD8^+^ T-cell hyporesponsiveness and increased Vδ1/Vδ2 γδ T-cell ratio – a surrogate marker for operational tolerance.^43^ Accordingly, the higher rate of asymptomatic CMV (re)activation found in the pre-emptive strategy group could potentially lead to regulatory γδ T-cell-based graft protection and explain the better graft function and lower incidence of acute rejection; an early administration of (val)ganciclovir would therefore hinder the development of this protective immune response. Our hypothesis is compatible with the observed differences between male and female recipients, as sex-associated differences in the anti-CMV immunity have been shown by Lindemann *et al.* to correlate with graft function.^13^ Even though this effect cannot explain our observations, it demonstrates how sex and anti-CMV immunity can potentially interact and affect eGFR.^13^ Therefore, further research, including systems medicine approaches, is needed to better understand the effects of CMV prevention strategies from an immune, virological and pharmacokinetic point of view – with emphasis on sex-associated differences – and their effects on transplantation outcome.^44,45^

Of interest, the prophylactic strategy group showed a higher incidence of late-onset CMV (from month six on); such increases of viral (re)activation incidence after the end of prophylaxis have been observed before.^30,31^ We also observed a tendency towards higher incidence of rejection and severe BK virus (re)activation among patients in the prophylactic strategy group, although these tendencies could not be confirmed through multi-parameter analysis. While we have already reported a negative effect of prophylaxis on rejection within the VIPP study – albeit only for the D^-^R^+^ subgroup – this study is the first to suggest such an association in the entire cohort.^23,24^ Regarding BKV, the observed pattern is in line with three recent studies.^46–48^ On the other hand, we did not observe any effect of prevention strategy on EBV (re)activation.^49^ This is relevant, as there is currently no consensus in the literature on this topic. Even though a number of publications have observed an effect against EBV or post-transplant lymphoproliferative disease (the main EBV-associated complication), a meta-study with 2366 participants saw no effect of prophylaxis for this EBV complication.^50–52^

This study is based on the prospective Harmony study, a trial designed with the goal of identifying which immunosuppressive drug combination is superior with respect to acute rejections and secondary to a number of other outcome variables, including graft function and viral (re)activations.^19^ A potential shortcoming of the present study consists therefore in the fact that prevention strategy groups were not randomized, and no power calculation was performed with respect to this question. Therefore, even though we have controlled for all measured demographic factors in the analyses and other potential confounding factors – including the first measured eGFR after transplantation – we cannot completely exclude bias in unmeasured factors as the cause of the observed differences. A further limitation is related to the criteria employed for deciding the prevention strategy for each patient: As the decision to adopt a prophylactic or a pre-emptive strategy was taken by each individual physician or centre, it is difficult to ascertain the background, potentially introducing bias in the use of prevention strategies. On the other hand, our study does have some advantages: We have analysed a larger (N=540) and more heterogeneous cohort (patients with all CMV mismatch-based risk constellations) than most studies on the matter, thereby achieving higher statistical power.^23,24,28–32^ Moreover, our study design is closer to clinical reality, with similar valganciclovir doses and prophylaxis duration to those routinely employed in the clinic.^53^ Based on the limitations and advantages of the study, we deem our results as evidence that further research is needed to determine the effects of prevention strategies on transplantation outcome and their hypothetical interactions with sex.

In summary, our study provides the first evidence in the literature suggesting superiority of the pre-emptive approach in female patients. Even though the prophylactic strategy was associated with reduced prevalence of CMV (re)activation and syndrome, it was associated with a strong impairment of the renal function. The effects of prevention strategy on graft function were shown in the multi-parameter analysis as independent from potential bias in the cohort. Moreover, we observed tentative evidence of a sex-independent tendency towards higher incidence of acute rejection and BKV (re)activation. Further randomized controlled studies are needed to confirm these observations.

## Authors’ Contributions

NB, NW, CH, OT, CB, KW, RS, OW, THW, BS, PR, and MO contributed to the study design, sample collection, and/or sample management. CDH carried out experiments. ABN, NB, and MO performed data interpretation. ABN, NB and MO drafted the manuscript. All authors have contributed to the manuscript and approved the final version of the manuscript for submission.

## Description of Supporting Information

Table S1. Differences in cause of end-stage kidney disease between strategy groups.

Table S2. Detailed results of the multi-parameter models of transplantation outcome.

Table S3. Differences in patient baseline characteristics between strategy groups within the female sub-cohort.

Figure S4. Box plot of the graft function dynamics of the prevention strategy groups. The numbers indicate the P value of the difference in eGFR between the prevention strategy groups, as calculated using the Mann-Whitney test. The P values for the first six measurements are included only to facilitate understanding on the eGFR dynamics, and are therefore not adjusted for multiple testing.

Table S5. Results of the single-parameter analyses for virus-related complications. The detailed results of single-parameter associations of prevention strategy, including stratified analysis for risk constellation. Following complications were analysed: CMV (re)activation and syndrome, EBV (re)activation and BKV (re)activation.

## References

1. Elfadawy N, Flechner SM, Liu X, Schold J, Srinivas TR, Poggio E, et al. CMV Viremia is associated with a decreased incidence of BKV reactivation after kidney and kidney-pancreas transplantation. Transplantation. 2013;96(12):1097–1103.

2. Fehr T, Cippà PE, Mueller NJ. Cytomegalovirus post kidney transplantation: Prophylaxis versus pre-emptive therapy? Transpl Int. 2015;28:1351–1356.

3. Le Page AK, Mackie FE, McTaggart SJ, Kennedy SE. Cytomegalovirus & Epstein Barr Virus serostatus as a predictor of the long-term outcome of kidney transplantation. Nephrology (Carlton). 2013;18(12):813–819.

4. Egli A, Binggeli S, Bodaghi S, Dumoulin A, Funk GA, Khanna N, et al. Cytomegalovirus and polyomavirus BK posttransplant. Nephrol Dial Transplant. 2007;22([Suppl 8]):viii72–viii82.

5. Malvezzi P, Jouve T, Rostaing L. Induction by anti-thymocyte globulins in kidney transplantation: a review of the literature and current usage. J Nephropathol. 2015;4(4):110–115.

6. Kotton CN, Kumar D, Caliendo AM, Huprikar S, Chou S, Danziger-Isakov L, et al. The Third International Consensus Guidelines on the Management of Cytomegalovirus in Solid-Organ Transplantation. Vol 102.; 2018.

7. Special Issue: KDIGO Clinical Practice Guideline for the Care of Kidney Transplant Recipients. Am J Transplant. 2009;9(Suppl 3):S1–S157.

8. Egli A, Binet I, Binggeli S, Jäger C, Dumoulin A, Schaub S, et al. Cytomegalovirus-specific T-cell responses and viral replication in kidney transplant recipients. J Transl Med. 2008;6:29.

9. Bunde T, Kirchner A, Hoffmeister B, Habedank D, Hetzer R, Cherepnev G, et al. Protection from cytomegalovirus after transplantation is correlated with immediate early 1–specific CD8 T cells. J Exp Med. 2005;201(7):1031–1036.

10. Sester M, Sester U, Gartner B, Heine G, Girndt M, Mueller-Lantzsch N, et al. Levels of virus-specific CD4 T cells correlate with cytomegalovirus control and predict virus-induced disease after renal transplantation. Transplantation. 2001;71(9):1287–1294.

11. Momper JD, Misel ML, McKay DB. Sex differences in transplantation. Transplant Rev. 2017;31(3):145–150.

12. Perrottet N, Csajka C, Pascual M, Manuel O, Lamoth F, Meylan P, et al. Population pharmacokinetics of ganciclovir in solid-organ transplant recipients receiving oral valganciclovir. Antimicrob Agents Chemother. 2009;53(7):3017–3023.

13. Lindemann M, Korth J, Sun M, Xu S, Struve C, Werner K, et al. The Cytomegalovirus-Specific IL-21 ELISpot Correlates with Allograft Function of Kidney Transplant Recipients. Int J Mol Sci. 2018;19(12):3945.

14. Villacres MC, Longmate J, Auge C, Diamond DJ. Predominant type 1 CMV-specific memory T-helper response in humans: Evidence for gender differences in cytokine secretion. Hum Immunol. 2004;65(5):476–485.

15. Fleck-Derderian S, McClellan W, Wojcicki JM. The association between cytomegalovirus infection, obesity, and metabolic syndrome in U.S. adult females. Obesity. 2017;25(3):626–633.

16. Hassan J, O’Neill D, Honari B, De Gascun C, Connell J, Keogan M, et al. Cytomegalovirus Infection in Ireland. Med. 2016;95(6):e2735.

17. Stranzinger J, Kindel J, Henning M, Wendeler D, Nienhaus A. Prevalence of CMV infection among staff in a metropolitan children’s hospital - occupational health screening findings. GMS Hyg Infect Control. 2016;11:Doc20.

18. Di Benedetto S, Derhovanessian E, Steinhagen-Thiessen E, Goldeck D, Müller L, Pawelec G. Impact of age, sex and CMV-infection on peripheral T cell phenotypes: results from the Berlin BASE-II Study. Biogerontology. 2015;16(5):631–643.

19. Thomusch O, Wiesener M, Opgenoorth M, Pascher A, Woitas RP, Witzke O, et al. Rabbit-ATG or basiliximab induction for rapid steroid withdrawal after renal transplantation (Harmony): an open-label, multicentre, randomised controlled trial. Lancet. 2016;388:3006–3016.

20. Blazquez-Navarro A, Dang-Heine C, Wittenbrink N, Bauer C, Wolk K, Sabat R, et al. BKV, CMV, and EBV Interactions and their Effect on Graft Function One Year Post-Renal Transplantation: Results from a Large Multi-Centre Study. EBioMedicine. 2018;34:113–121.

21. Levey AS, Stevens LA, Schmid CH, Zhang YL, Castro AF, Feldman HI, et al. A new equation to estimate glomerular filtration rate. Ann Intern Med. 2009;150(9):604–612.

22. Solez K, Colvin RB, Racusen LC, Sis B, Halloran PF, Birk PE, et al. Banff ‘05 meeting report: Differential diagnosis of chronic allograft injury and elimination of chronic allograft nephropathy (‘CAN’). Am J Transplant. 2007;7(3):518–526.

23. Witzke O, Nitschke M, Bartels M, Wolters H, Wolf G, Reinke P, et al. Valganciclovir prophylaxis versus preemptive therapy in cytomegalovirus-positive renal allograft recipients: Long-term results after 7 years of a randomized clinical trial. Transplantation. 2018;102:876–882.

24. Witzke O, Hauser IA, Bartels M, Wolf G, Wolters H, Nitschke M. Valganciclovir Prophylaxis Versus Preemptive Therapy in Cytomegalovirus-Positive Renal Allograft Recipients: 1-Year Results of a Randomized Clinical Trial. Transplantation. 2012;93:61–68.

25. Velentgas P, Dreyer NA, Nourjah P, Smith SR, Torchia MM, eds. Developing a Protocol for Observational Comparative Effectiveness Research A User’s Guide. Rockville, MD: Agency for Healthcare Research and Quality; 2013.

26. Bender R, Lange S. Adjusting for multiple testing - when and how? J Clin Epidemiol. 2001;54:343–349.

27. Li G, Taljaard M, Van den Heuvel ER, Levine MA, Cook DJ, Wells GA, et al. An introduction to multiplicity issues in clinical trials: the what, why, when and how. Int J Epidemiol. 2017:746–755.

28. Kliem V, Fricke L, Wollbrink T, Burg M, Radermacher J, Rohde F. Improvement in long-term renal graft survival due to CMV prophylaxis with oral ganciclovir: Results of a randomized clinical trial. Am J Transplant. 2008;8(5):975–983.

29. Spinner ML, Saab G, Casabar E, Bowman LJ, Gregory A, Brennan DC. Impact of Prophylactic Versus Preemptive Valganciclovir on Long-term Renal Allograft Outcomes. Transplantation. 2010;90(4):412–418.

30. Florescu DF, Qiu F, Schmidt CM, Kalil AC. A direct and indirect comparison meta-analysis on the efficacy of cytomegalovirus preventive strategies in solid organ transplant. Clin Infect Dis. 2014;58(6):785–803.

31. Khoury JA, Storch GA, Bohl DL, Schuessler RM, Torrence SM, Lockwood M, et al. Prophylactic versus preemptive oral valganciclovir for the management of cytomegalovirus infection in adult renal transplant recipients. Am J Transplant. 2006;6(9):2134–2143.

32. Van Der Beek MT, Berger SP, Vossen ACTM, Van Der Blij-De Brouwer CS, Press RR, De Fijter JW, et al. Preemptive versus sequential prophylactic-preemptive treatment regimens for cytomegalovirus in renal transplantation: Comparison of treatment failure and antiviral resistance. Transplantation. 2010;89(3):320–326.

33. Small LN, Lau J, Snydman DR. Preventing Post–Organ Transplantation Cytomegalovirus Disease with Ganciclovir: A Meta-Analysis Comparing Prophylactic and Preemptive Therapies. Clin Infect Dis. 2005;43(May):869–880.

34. Meije Y, Fortún J, Len O, Aguado JM, Moreno A, Cisneros JM, et al. Prevention strategies for cytomegalovirus disease and long-term outcomes in the high-risk transplant patient (D+/R-): Experience from the RESITRA-REIPI cohort. Transpl Infect Dis. 2014;16(3):387–396.

35. Kasiske BL, Israni AK, Snyder JJ, Skeans MA. The relationship between kidney function and long-term graft survival after kidney transplant. Am J Kidney Dis. 2011;57(3):466–475.

36. Rásky É, Waxenegger A, Groth S, Stolz E, Schenouda M, Berzlanovich A. Sex and gender matters: A sex-specific analysis of original articles published in the Wiener klinische Wochenschrift between 2013 and 2015. Wien Klin Wochenschr. 2017;129:781–785.

37. Meier-Kriesche HU, Ojo AO, Leavey SF, Hanson JA, Leichtman AB, Magee JC, et al. Gender differences in the risk for chronic renal allograft failure. Transplantation. 2001;71(3):429–432.

38. Hirsch HH, Vincenti F, Friman S, Tuncer M, Citterio F, Wiecek A, et al. Polyomavirus BK replication in de novo kidney transplant patients receiving tacrolimus or cyclosporine: A prospective, randomized, multicenter study. Am J Transplant. 2013;13(1):136–145.

39. Salvadori M, Rosati A, Bock A, Chapman J, Dussol B, Fritsche L, et al. Estimated one-year glomerular filtration rate is the best predictor of long-term graft function following renal transplant. Transplantation. 2006;81(2):202–206.

40. Schwarz A, Linnenweber-Held S, Heim A, Bröcker V, Rieck D, Framke T, et al. Factors influencing viral clearing and renal function during polyomavirus BK-associated nephropathy after renal transplantation. Transplantation. 2012;94(4):396–402.

41. Heldenbrand S, Li C, Cross RP, DePiero KA, Dick TB, Ferguson K, et al. Multicenter evaluation of efficacy and safety of low-dose versus high-dose valganciclovir for prevention of cytomegalovirus disease in donor and recipient positive (D+/R+) renal transplant recipients. Transpl Infect Dis. 2016;18(6):904–912.

42. Nickel P, Bold G, Presber F, Biti D, Babel N, Kreutzer S, et al. High levels of CMV-IE-1-specific memory T cells are associated with less alloimmunity and improved renal allograft function. Transpl Immunol. 2009;20:238–242.

43. Shi XL, De Mare-Bredemeijer ELD, Tapirdamaz Ö, Hansen BE, Van Gent R, Van Campenhout MJH, et al. CMV Primary Infection Is Associated with Donor-Specific T Cell Hyporesponsiveness and Fewer Late Acute Rejections after Liver Transplantation. Am J Transplant. 2015;15:2431–2442.

44. Blazquez-Navarro A, Schachtner T, Stervbo U, Sefrin A, Stein M, Westhoff TH, et al. Differential T cell response against BK virus regulatory and structural antigens: A viral dynamics modelling approach. PLOS Comput Biol. 2018;14(5):e1005998.

45. Maier M, Takano T, Sapir-Pichhadze R. Changing paradigms in the management of rejection in kidney transplantation: Evolving from protocol-based care to the era of P4 medicine. Can J Kidney Heal Dis. 2017;4:1–12.

46. Reischig T, Kacer M, Hruba P, Hermanova H, Hes O, Lysak D, et al. Less renal allograft fibrosis with valganciclovir prophylaxis for cytomegalovirus compared to high-dose valacyclovir: A parallel group, open-label, randomized controlled trial 11 Medical and Health Sciences 1103 Clinical Sciences. BMC Infect Dis. 2018;18(1):1–9.

47. Reischig T, Kacer M, Jindra P, Hes O, Lysak D, Bouda M. Randomized trial of valganciclovir versus valacyclovir prophylaxis for prevention of cytomegalovirus in renal transplantation. Clin J Am Soc Nephrol. 2015;10(2):294–304.

48. Reischig T, Kacer M, Hes O, Machova J, Nemcova J, Lysak D, et al. Cytomegalovirus prevention strategies and the risk of BK polyomavirus viremia and nephropathy. Am J Transplant. 2019.

49. Végso G, Hajdu M, Sebestyén A. Lymphoproliferative disorders after solid organ transplantation-classification, incidence, risk factors, early detection and treatment options. Pathol Oncol Res. 2011;17(3):443–454.

50. Cameron BM, Kennedy SE, Rawlinson WD, Mackie FE. The efficacy of valganciclovir for prevention of infections with cytomegalovirus and Epstein-Barr virus after kidney transplant in children. Pediatr Transplant. 2017;21(1):1–11.

51. Funch DP, Walker AM, Schneider G, Ziyadeh NJ, Pescovitz MD. Ganciclovir and acyclovir reduce the risk of post-transplant lymphoproliferative disorder in renal transplant recipients. Am J Transplant. 2005;5(12):2894–2900.

52. Aldabbagh MA, Gitman MR, Kumar D, Humar A, Rotstein C, Husain S. The Role of Antiviral Prophylaxis for the Prevention of Epstein-Barr Virus-Associated Posttransplant Lymphoproliferative Disease in Solid Organ Transplant Recipients: A Systematic Review. Am J Transplant. 2016:770–781.

53. Rissling O, Naik M, Brakemeier S, Schmidt D, Staeck O, Hohberger A, et al. High frequency of valganciclovir underdosing for cytomegalovirus prophylaxis after renal transplantation. Clin Kidney J. 2018;11(4):564–573.

