## Supplementary material for "Sex-associated differences in cytomegalovirus prevention: Prophylactic strategy is associated with a strong kidney function impairment in female renal transplant patients": Table S1

| Cause of end-stage kidney disease | Prophylactic strategy group (N=308) | Pre-emptive strategy group (N=232) | P Value |
|---|---|---|---|
| Hypertension or large vessel disease | 111 (36.0%) | 87 (37.8%) | 0.738 |
| Glomerulonephritis | 79 (25.6%) | 69 (30.0%) | 0.308 |
| Polycystic kidney disease (adult type, dominant) | 64 (20.8%) | 38 (16.5%) | 0.256 |
| Diabetes | 32 (10.4%) | 23 (10.0%) | 0.997 |
| Interstitial nephritis or pyelonephritis | 20 (6.5%) | 19 (8.3%) | 0.539 |
| Secondary glomerulonephritis or vasculitis | 7 (2.3%) | 4 (1.7%) | 0.765[a] |
| Other hereditary or congenital diseases | 11 (3.6%) | 8 (3.5%) | 1.000 |
| Neoplasms or tumours | 4 (1.3%) | 0 (0.0%) | 0.139[a] |
| Other | 109 (35.4%) | 72 (31.3%) | 0.368 |
| Undefined cause | 27 (9.3%) | 24 (11.0%) | 0.621 |

**Table S1 – Differences in cause of end-stage kidney disease between strategy groups.** Data are given in number (percentage). P value is calculated based on Pearson's chi-square test or Fisher's exact test (marked with [a]). Causes of end-stage kidney disease are not mutually exclusive.
