## Supplementary material for "Sex-associated differences in cytomegalovirus prevention: Prophylactic strategy is associated with a strong kidney function impairment in female renal transplant patients": Table S2

| Explanatory variables | Estimate | Standard error | P value |
|---|---|---|---|
| (Intercept) | 102.7485 | 7.5093 | <0.001 |
| Prophylactic strategy | -1.0719 | 2.4167 | 0.658 |
| Female sex | 4.6257 | 3.1683 | 0.146 |
| Prophylactic strategy:Female sex | -12.0421 | 4.2285 | 0.005 |
| Recipient age (years) | -0.1879 | 0.0925 | 0.043 |
| Donor age (years) | -0.4339 | 0.0719 | <0.001 |
| Body mass index ($kg \cdot m^{-2}$) | -0.7971 | 0.2156 | <0.001 |
| No panel-reactive antibodies before transplantation | -4.9624 | 3.5436 | 0.163 |
| Cause of end-stage renal disease: Diabetes | 5.8559 | 3.7196 | 0.117 |
| Tacrolimus trough level ($ng \cdot mL^{-1}$) | 0.4209 | 0.2330 | 0.072 |
| Acute rejection | -8.2117 | 3.3308 | 0.014 |
| Peak BKV viral load | -0.9365 | 0.5187 | 0.072 |

**Table S2 A – Multivariate analysis of prevention strategy and sex effects on eGFR-1y.** eGFR-1y was estimated by linear regression. Confounders were selected employing backwards elimination, starting with all demographic factors (Table 1 and Table S1), CMV, BKV and EBV peak viral loads, acute rejection and transplantation centre.

BK virus (BKV), Estimated glomerular filtration rate one year after transplantation (eGFR-1y)

| Explanatory variables | Estimate | Standard error | P value |
|---|---|---|---|
| (Intercept) | 65.8128 | 4.8610 | <0.001 |
| Prophylactic strategy | -0.9832 | 2.0679 | 0.635 |
| Female sex | 3.5030 | 2.6976 | 0.195 |
| Prophylactic strategy:Female sex | -10.2195 | 3.5922 | 0.005 |
| eGFR-2w | 0.4613 | 0.0434 | <0.001 |
| Recipient age (years) | -0.1993 | 0.0791 | 0.012 |
| Donor age (years) | -0.3597 | 0.0618 | <0.001 |
| Cause of end-stage renal disease: Diabetes | 5.2122 | 3.1380 | 0.098 |
| White blood cell count (cells·$L^{-1}$) | -0.0027 | 0.0018 | 0.143 |
| Acute rejection | -5.2399 | 2.8616 | 0.068 |
| Peak BKV viral load | -0.8926 | 0.4444 | 0.046 |

**Table S2 B – Multivariate analysis of prevention strategy and sex effects on eGFR-1y controlling for eGFR-2w.** eGFR-1y was estimated by linear regression. Confounders were selected employing backwards elimination, starting with all demographic factors (Table 1 and Table S1), CMV, BKV and EBV peak viral loads, acute rejection and transplantation centre.

BK virus (BKV), Estimated glomerular filtration rate two weeks after transplantation (eGFR-2w), Estimated glomerular filtration rate one year after transplantation (eGFR-1y)

| Explanatory variables | Estimate | Standard error | P value |
|---|---|---|---|
| (Intercept) | -4.7551 | 0.6773 | <0.001 |
| Prophylactic strategy | 0.9161 | 0.5021 | 0.068 |
| Female sex | 0.3380 | 0.6369 | 0.596 |
| Prophylactic strategy:Female sex | -0.7495 | 0.7708 | 0.331 |
| Number of HLA A, B and DR mismatches | 0.3724 | 0.1179 | 0.002 |
| Donor with expanded criteria | 0.5516 | 0.3738 | 0.140 |
| Cause of end-stage renal disease: Polycystic kidney disease (adult type, dominant) | 1.1651 | 0.4545 | 0.010 |
| Cause of end-stage renal disease: Interstitial nephritis or pyelonephritis | 1.2149 | 0.5925 | 0.040 |
| Cause of end-stage renal disease: Other | 1.0503 | 0.4050 | 0.010 |

**Table S2 C – Multivariate analysis of prevention strategy and sex effects on acute rejection.** Occurrence of acute rejection during the first post-transplantation year was estimated by logistic regression. Confounders were selected employing backwards elimination, starting with all demographic factors (Table 1 and Table S1) and transplantation centre.

| Explanatory variables | Estimate | Standard error | P value |
|---|---|---|---|
| (Intercept) | 1.3471 | 0.2842 | <0.001 |
| Prophylactic strategy | -0.7348 | 0.2339 | 0.002 |
| Female sex | -0.0434 | 0.2917 | 0.882 |
| Prophylactic strategy:Female sex | -0.0820 | 0.3593 | 0.820 |
| CMV mismatch-based risk: Medium ($R^+$) | -0.3314 | 0.2187 | 0.131 |
| CMV mismatch-based risk: Low ($D^-R^-$) | -1.2084 | 0.2215 | <0.001 |
| Cause of end-stage renal disease: Other hereditary or congenital diseases | 1.3444 | 0.6633 | 0.044 |
| White blood cell count (cells·$L^{-1}$) | 0.0004 | 0.0086 | 0.967 |
| Therapy arm: arm B (basiliximab) | 0.1152 | 0.1866 | 0.537 |
| Therapy arm: arm C (ATG) | 0.4704 | 0.2046 | 0.022 |

**Table S2 D – Multivariate analysis of prevention strategy and sex effects on CMV peak viral load.** Peak viral load in logarithmic scale (with a value of 0 for viral load below detection limit) was estimated by linear regression. Confounders were selected employing backwards elimination, starting with all demographic factors (Table 1 and Table S1) and transplantation centre.

Antithymocyte globulin (ATG), Cytomegalovirus (CMV), Seronegative donor and seronegative recipient ($D^-R^-$), Seropositive donor and seronegative recipient ($D^+R^-$), Mycophenolate mofetil (MMF), Seropositive Recipient ($R^+$)

| Explanatory variables | Estimate | Standard error | P value |
|---|---|---|---|
| (Intercept) | 0.4760 | 0.8678 | 0.583 |
| Prophylactic strategy | -1.1650 | 0.5734 | 0.042 |
| Female sex | -0.3126 | 0.6749 | 0.643 |
| Prophylactic strategy:Female sex | 0.8465 | 0.8551 | 0.322 |
| Donor age (years) | 0.0332 | 0.0131 | 0.011 |
| CMV mismatch-based risk: Medium ($R^+$) | -0.9913 | 0.4585 | 0.031 |
| CMV mismatch-based risk: Low ($D^-R^-$) | -4.9670 | 1.2380 | <0.001 |
| Living donor | -2.9140 | 0.8710 | 0.001 |
| Cause of end-stage renal disease: Polycystic kidney disease (adult type, dominant) | -0.8544 | 0.4954 | 0.085 |
| Cause of end-stage renal disease: Diabetes | 1.4870 | 0.6509 | 0.022 |
| Cause of end-stage renal disease: Undefined cause | -1.6880 | 0.7436 | 0.023 |
| White blood cell count (cells·$L^{-1}$) | 0.0022 | 0.0081 | 0.786 |
| Centre effects | - | - | <0.001 |

**Table S2 C – Multivariate analysis of prevention strategy and sex effects on CMV syndrome.** Occurrence of CMV syndrome during the first post-transplantation year was estimated by logistic regression. Confounders were selected employing backwards elimination, starting with all demographic factors (Table 1 and Table S1) and transplantation centre. The P value for centre effects refers to the minimum P value of any transplantation centre.

Cytomegalovirus (CMV), Seronegative donor and seronegative recipient ($D^-R^-$), Seropositive donor and seronegative recipient ($D^+R^-$), Epstein-Barr virus (EBV), Seropositive Recipient ($R^+$)

| Explanatory variables | Estimate | Standard error | P value |
|---|---|---|---|
| (Intercept) | 1.1206 | 0.3928 | 0.005 |
| Prophylactic strategy | 0.0713 | 0.1990 | 0.720 |
| Female sex | 0.1616 | 0.2587 | 0.533 |
| Prophylactic strategy:Female sex | -0.3769 | 0.3419 | 0.271 |
| Number of HLA A, B and DR mismatches | 0.1008 | 0.0511 | 0.050 |
| Donor with expanded criteria | 0.3255 | 0.1732 | 0.061 |
| No panel-reactive antibodies before transplantation | -0.5790 | 0.2908 | 0.048 |
| Cause of end-stage renal disease: Hypertension | 0.2967 | 0.1755 | 0.092 |
| Cause of end-stage renal disease: Polycystic kidney disease (adult type, dominant) | -0.5602 | 0.2158 | 0.010 |
| Cause of end-stage renal disease: Diabetes | -0.5898 | 0.2944 | 0.046 |
| Cause of end-stage renal disease: Neoplasms or tumours | 2.6395 | 0.9303 | 0.005 |
| Cause of end-stage renal disease: Other | -0.2504 | 0.1775 | 0.160 |
| Cause of end-stage renal disease: Undefined cause | -0.4438 | 0.2682 | 0.099 |
| Low MMF daily dose (< 2000 $mg \cdot day^{-1}$) | -0.0003 | 0.0002 | 0.083 |

**Table S2 F – Multivariate analysis of prevention strategy and sex effects on EBV peak viral load.** Peak viral load in logarithmic scale (with a value of 0 for viral load below detection limit) estimated by linear regression. Confounders were selected employing backwards elimination, starting with all demographic factors (Table 1 and Table S1) and transplantation centre.

Epstein-Barr virus (EBV), Mycophenolate mofetil (MMF)

| Explanatory variables | Estimate | Standard error | P value |
|---|---|---|---|
| (Intercept) | 1.8460 | 0.2877 | <0.001 |
| Prophylactic strategy | 0.5597 | 0.2907 | 0.055 |
| Female sex | -0.2151 | 0.3476 | 0.537 |
| Prophylactic strategy:Female sex | -0.3282 | 0.4734 | 0.489 |
| Number of HLA A, B and DR mismatches | -0.1046 | 0.0711 | 0.143 |
| Cause of end-stage renal disease: Glomerulonephritis | -0.3842 | 0.2776 | 0.168 |
| Cause of end-stage renal disease: Interstitial nephritis or pyelonephritis | 0.7469 | 0.4619 | 0.107 |
| Cause of end-stage renal disease: Other | 0.4241 | 0.2724 | 0.121 |

**Table S2 G – Multivariate analysis of prevention strategy and sex effects on BKV peak viral load.** Peak viral load in logarithmic scale (with a value of 0 for viral load below detection limit) estimated by linear regression. Confounders were selected employing backwards elimination, starting with all demographic factors (Table 1 and Table S1) and transplantation centre.

BK virus (BKV)
