## Supplementary material for "Sex-associated differences in cytomegalovirus prevention: Prophylactic strategy is associated with a strong kidney function impairment in female renal transplant patients": Table S3

| Variable | | Females in prophylactic strategy group (N=104) | Females in pre-emptive strategy group (N=90) | P value |
|---|---|---|---|---|
| Caucasian race | | 104 (100.0%) | 90 (100.0%) | - |
| Recipient age (years) | | 57 [47-65] | 58.5 [48.2-64] | 0.910 |
| Body mass index ($kg \cdot m^{-2}$) | | 26.5 [23.5-30.9] | 24.5 [21.9-27.9] | 0.005 |
| CMV mismatch -based risk | High ($D^+R^-$) | 36 (35%) | 8 (9.3%) | <0.001 |
| | Medium ($R^+$) | 55 (53.4%) | 55 (64%) | |
| | Low ($D^-R^-$) | 12 (11.7%) | 23 (26.7%) | |
| EBV mismatch -based risk | High ($D^+R^-$) | 2 (2.4%) | 3 (4.4%) | 0.373[a] |
| | Medium ($R^+$) | 82 (96.5%) | 62 (91.2%) | |
| | Low ($D^-R^-$) | 1 (1.2%) | 3 (4.4%) | |
| Donor age (years) | | 58.5 [48-69] | 56 [45-64] | 0.183 |
| No previous transplantations | | 101 (97.1%) | 85 (95.5%) | 0.705[a] |
| Living donor | | 14 (13.5%) | 15 (17%) | 0.625 |
| Expanded criteria donor | | 57 (54.8%) | 37 (41.1%) | 0.078 |
| Cold ischaemia time (min) | | 620.5 [419-881.8] | 656 [421-862.5] | 0.949 |
| Number of HLA A, B and DR mismatches | | 3 [2-4] | 3 [1-4] | 0.087 |
| No panel-reactive antibodies before transplantation | | 12 (11.8%) | 9 (10.5%) | 0.961 |
| White blood cell count ($cells \cdot L^{-1}$) | | 7.5 [6-9.2] | 7.2 [6.1-9.3] | 0.655 |
| Therapy arm | A (basiliximab+steroids) | 28 (26.9%) | 35 (38.9%) | 0.106 |
| | B (basiliximab) | 35 (33.7%) | 31 (34.4%) | |
| | C (ATG) | 41 (39.4%) | 24 (26.7%) | |

| | | | | |
|---|---|---|---|---|
| Low MMF daily dose (< 2000 mg·day$^{-1}$) | | 17 (16.3%) | 16 (17.8%) | 0.942 |
| Tacrolimus trough level (ng·mL$^{-1}$) | | 9.9 [7.4-13.2] | 8.9 [6.9-11.2] | 0.130 |
| Cause of end-stage kidney disease | Hypertension or large vessel disease | 31 (29.8%) | 35 (38.9%) | 0.238 |
| | Glomerulonephritis | 19 (18.3%) | 26 (28.9%) | 0.115 |
| | Polycystic kidney disease (adult type, dominant) | 29 (27.9%) | 14 (15.6%) | 0.059 |
| | Diabetes | 11 (10.6%) | 4 (4.4%) | 0.185 |
| | Interstitial nephritis or pyelonephritis | 11 (10.6%) | 14 (15.6%) | 0.414 |
| | Secondary glomerulonephritis or vasculitis | 2 (1.9%) | 2 (2.2%) | 1.000[a] |
| | Other hereditary or congenital diseases | 6 (5.8%) | 3 (3.3%) | 0.508[a] |
| | Neoplasms or tumours | 0 (0.0%) | 0 (0.0%) | - |
| | Other | 31 (29.8%) | 29 (32.2%) | 0.836 |
| | Undefined cause | 9 (9.3%) | 6 (7.1%) | 0.803 |

**Table S3 – Differences in patient baseline characteristics between strategy groups within the female sub-cohort.** Data are given as number (percentage) or median [interquartile range]. Expanded criteria donors are defined as follows: age over 60 years or age over 50 years and at least two of the following factors: cerebrovascular accident as the cause of death, hypertension or a serum creatinine level over 1.5 mg·dL$^{-1}$. P value is calculated based on Pearson's chi-square test or Fisher's exact test for binary variables (marked with [a]) and based on Mann-Whitney test for continuous variables.
