## Supplementary figures and images for "Sex-associated differences in cytomegalovirus prevention: Prophylactic strategy is associated with a strong kidney function impairment in female renal transplant patients"

### Figure S4

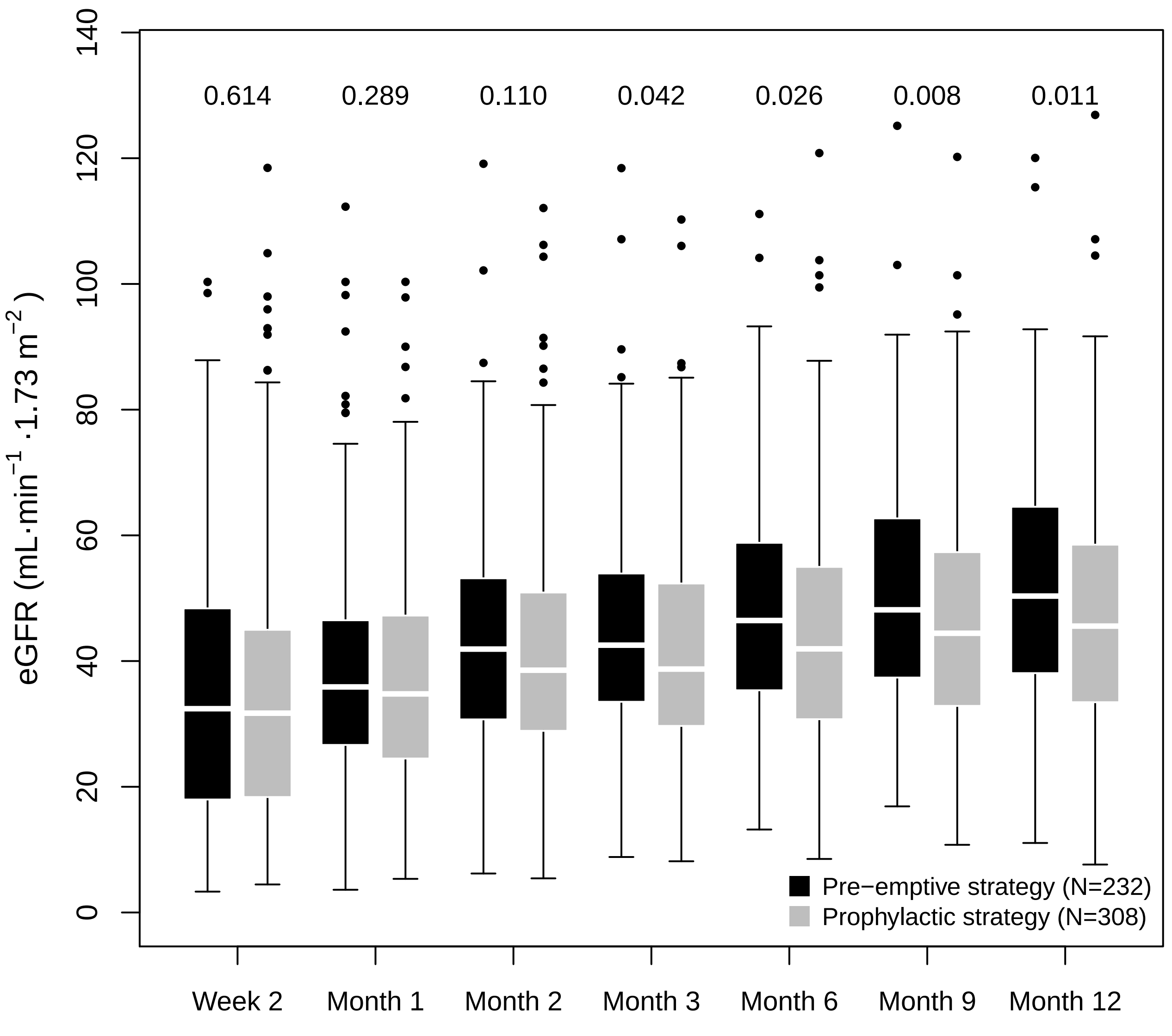
