## Supplementary material for "Sex-associated differences in cytomegalovirus prevention: Prophylactic strategy is associated with a strong kidney function impairment in female renal transplant patients": Table S5

| Viral Event | Severity | Prophylactic strategy group (N=308) | Pre-emptive strategy group (N=232) | P value | OR [95% CI] |
|---|---|---|---|---|---|
| CMV (re)activation | Detectable viral load | 46 (14.9%) | 46 (19.8%) | 0.167 | 1.41 [0.87-2.27] |
| | Elevated viral load | 19 (6.2%) | 20 (8.6%) | 0.357 | 1.43 [0.71-2.92] |
| | High viral load | 10 (3.2%) | 8 (3.4%) | 1.000 | 1.06 [0.36-3.05] |
| CMV syndrome | All | 78 (25.3%) | 35 (15.1%) | 0.005 | 0.52 [0.33-0.83] |
| | Mild | 39 (50.0%) | 24 (68.6%) | 0.204 | - |
| | Moderate | 31 (39.7%) | 9 (25.7%) | | |
| | Severe | 8 (10.3%) | 2 (5.7%) | | |
| EBV (re)activation | Detectable viral load | 62 (20.1%) | 47 (20.3%) | 1.000 | 1.01 [0.64-1.57] |
| | Elevated viral load | 17 (5.5%) | 20 (8.6%) | 0.215 | 1.61 [0.78-3.37] |
| | High viral load | 6 (1.9%) | 5 (2.2%) | 1.000[a] | 1.11 [0.26-4.42] |
| BKV (re)activation | Detectable viral load | 157 (51.0%) | 103 (44.4%) | 0.153 | 0.77 [0.54-1.10] |
| | Elevated viral load | 78 (25.3%) | 43 (18.5%) | 0.077 | 0.67 [0.43-1.04] |
| | High viral load | 41 (13.3%) | 18 (7.8%) | 0.056 | 0.55 [0.29-1.01] |

**Table S5 A – Differences in viral events between strategy groups.** Data are given in number (percentage). P value is calculated based on Pearson's chi-square test or Fisher's exact test (marked with [a]). Odds ratio and confidence intervals are given only for viral events that were significant different between the strategy groups. For the definition of (re)activation severity degrees see Methods (2.7). With respect to severity of CMV syndrome, the percentage refers to the total number of CMV syndromes.

BK virus (BKV), Cytomegalovirus (CMV), 95% Confidence interval (95% CI), Epstein-Barr virus (EBV), Odds ratio (OR).

| CMV risk constellation | CMV event | Severity | Prophylactic strategy group (N=308) | Pre-emptive strategy group (N=232) | P value | OR [95% CI] |
|---|---|---|---|---|---|---|
| $D^+R^-$ (N=146) | (re)activation | Detectable | 31 (26.1%) | 9 (33.3%) | 0.598 | 1.42 [0.51-3.75] |
| | | Elevated | 13 (10.9%) | 6 (22.2%) | 0.123[a] | 2.31 [0.65-7.48] |
| | | High | 6 (5.0%) | 3 (11.1%) | 0.368[a] | 2.34 [0.35-11.9] |
| | Syndrome | All | 43 (36.1%) | 8 (29.6%) | 0.677 | 0.75 [0.26-1.97] |
| | | Mild | 14 (32.6%) | 6 (75.0%) | 0.062[a] | - |
| | | Moderate | 22 (51.2%) | 1 (12.5%) | | |
| | | Severe | 7 (16.3%) | 1 (12.5%) | | |
| $R^+$ (N=266) | (re)activation | Detectable | 12 (8.8%) | 35 (27.1%) | <0.001 | 3.86 [1.84-8.63] |
| | | Elevated | 5 (3.6%) | 13 (10.1%) | 0.066 | 2.95 [0.95-10.88] |
| | | High | 3 (2.2%) | 5 (3.9%) | 0.490[a] | 1.8 [0.34-11.81] |
| | Syndrome | All | 30 (21.9%) | 27 (20.9%) | 0.966 | 0.94 [0.5-1.77] |
| | | Mild | 23 (76.7%) | 18 (66.7%) | 0.460[a] | - |
| | | Moderate | 7 (23.3%) | 8 (29.6%) | | |
| | | Severe | 0 (0.0%) | 1 (3.7%) | | |
| $D^-R^-$ (N=115) | (re)activation | Detectable | 2 (4.2%) | 1 (1.5%) | 0.570[a] | 0.35 [0.01-6.94] |
| | | Elevated | 0 (0.0%) | 1 (1.5%) | 1.000[a] | - |
| | | High | 0 (0.0%) | 0 (0.0%) | - | - |
| | Syndrome | All | 3 (6.2%) | 0 (0.0%) | 0.070 | 0 [0-1.7] |
| | | Mild | 2 (66.7%) | 0 (0.0%) | 1.000[a] | - |
| | | Moderate | 0 (0.0%) | 0 (0.0%) | | |
| | | Severe | 1 (33.3%) | 0 (0.0%) | | |
| Arm C (N=176) | (re)activation | Detectable | 21 (17.1%) | 9 (17%) | 1.000 | 0.99 [0.37-2.49] |
| | | Elevated | 10 (8.1%) | 5 (9.4%) | 0.774[a] | 1.18 [0.3-4.02] |
| | | High | 5 (4.1%) | 2 (3.8%) | 1.000[a] | 0.93 [0.09-5.88] |
| | Syndrome | All | 29 (23.6%) | 8 (15.1%) | 0.287 | 0.58 [0.21-1.43] |
| | | Mild | 14 (48.3%) | 4 (50.0%) | 0.878[a] | - |
| | | Moderate | 10 (34.5%) | 2 (25.0%) | | |
| | | Severe | 5 (17.2%) | 2 (25.0%) | | |

**Table S5 B – Differences in cytomegalovirus complications between strategy groups stratified for CMV risk constellation.** Data are given in number (percentage). P value is calculated based on Pearson's chi-square test or Fisher's exact test (marked with [a]). Odds ratio and confidence intervals are given only for viral events that were significant different between the strategy groups. For the

definition of (re)activation severity degrees see Methods (2.7). With respect to severity of CMV syndrome, the percentage refers to the total number of CMV syndromes.
95% Confidence interval (95% CI), Cytomegalovirus (CMV), Seronegative donor and seronegative recipient ($D^-R^-$), Seropositive donor and seronegative recipient ($D^+R^-$), Odds ratio (OR), Seropositive Recipient ($R^+$).

| EBV risk constellation | Severity of (re)activation | Prophylactic strategy group (N=308) | Pre-emptive strategy group (N=232) | P value | OR [95% CI] |
|---|---|---|---|---|---|
| $D^+R^-$ (N=24) | Detectable | 3 (23.1%) | 3 (27.3%) | 1.000[a] | 1.24 [0.13-12] |
| | Elevated | 2 (15.4%) | 3 (27.3%) | 0.630[a] | 2.00 [0.18-29.33] |
| | High | 1 (7.7%) | 1 (9.1%) | 1.000[a] | 1.19 [0.01-101.86] |
| $R^+$ (N=400) | Detectable | 49 (20.5%) | 32 (19.9%) | 0.979 | 0.96 [0.56-1.63] |
| | Elevated | 12 (5.0%) | 14 (8.7%) | 0.209 | 1.80 [0.75-4.39] |
| | High | 5 (2.1%) | 3 (1.9%) | 1.000[a] | 0.89 [0.14-4.64] |
| $D^-R^-$ (N=9) | Detectable | 1 (25.0%) | 1 (20.0%) | 1.000[a] | 0.77 [0.01-78.24] |
| | Elevated | 0 (0.0%) | 0 (0.0%) | - | - |
| | High | 0 (0.0%) | 0 (0.0%) | - | - |
| Arm C (N=176) | Detectable | 28 (22.8%) | 18 (34%) | 0.173 | 1.74 [0.80-3.73] |
| | Elevated | 8 (6.5%) | 6 (11.3%) | 0.362[a] | 1.83 [0.49-6.39] |
| | High | 2 (1.6%) | 3 (5.7%) | 0.162[a] | 3.60 [0.40-44.31] |

**Table S5 C – Differences in EBV (re)activations between strategy groups stratified for EBV risk constellation.** Data are given in number (percentage). P value is calculated based on Pearson's chi-square test or Fisher's exact test (marked with [a]). Odds ratio and confidence intervals are given only for viral events that were significant different between the strategy groups. For the definition of (re)activation severity degrees, see Methods (2.7).
95% Confidence interval (95% CI), Seronegative donor and seronegative recipient ($D^-R^-$), Seropositive donor and seronegative recipient ($D^+R^-$), Epstein-Barr virus (EBV), Odds ratio (OR), Seropositive Recipient ($R^+$).
